# Atoh8 is a regulator of intestinal microfold cell (M cell) differentiation

**DOI:** 10.1101/2021.05.10.443378

**Authors:** Joel Johnson George, Laura Martin-Diaz, Markus Ojanen, Keijo Viiri

**Affiliations:** Faculty of Medicine and Health Technology, Tampere University Hospital, Tampere University Tampere, Finland

**Keywords:** Gut immunity, M cells, Peyer’s patch, Rank, RankL, Transcytosis

## Abstract

Intestinal microfold cells (M cells) are a dynamic lineage of epithelial cells that initiate mucosal immunity in the intestine. They are responsible for the uptake and transcytosis of microorganisms, pathogens and other antigens in the gastrointestinal tract. A mature M cell expresses a receptor Gp2 which binds to pathogens and aids in the uptake. Due to the rarity of these cells in the intestine, its development and differentiation remains yet to be fully understood. We recently demonstrated that polycomb repressive complex 2 (PRC2) is an epigenetic regulator of M cell development and 12 novel transcription factors including Atoh8 were revealed to be regulated by the PRC2. Here, we show that Atoh8 acts as a regulator of M cell differentiation; absence of Atoh8 led to a significant increase in the number of Gp2+ mature M cells and other M cell associated markers. Atoh8 null mice showed an increase in transcytosis capacity of luminal antigens. Increase in M cell population has been previously reported to be detrimental to mucosal immunity because some pathogens like orally acquired prions have been able to exploit the transcytosis capacity of M cells to infect the host; mouse with increased population of M cells are also susceptible to Salmonella infections. Our study here demonstrates that the population density of intestinal M-cell in the Peyer’s patch is regulated by the PRC2 regulated Atoh8.

## Introduction

The gastrointestinal tract is subject to constant exposure from antigens, microorganisms and foreign pathogens. The mucosal lining of the intestinal tract employs multiple mechanisms for immunosurveillance. These include epithelial tight junctions, production of antimicrobial peptide by Paneth cells, mucins from goblet cells, innate antigen receptors and acquired immunity in the form of secretory IgA (Liévin-Le Moal and Servin, 2006). Phagocytic and transcytosis capabilities are critical to induce an antigen-specific immune response. For this, the mucosal immune system is organized into inductive tissues such as the gut associated lymphoid tissue (GALT), these include a dome shaped specialized region known as Peyer’s patches (PP’s) in the intestine. PP’s are covered by dome-shaped follicle associated epithelium FAE which are composed of specialized intestinal epithelial cells (IEC’s) known as Microfold cells (M cells)(Owen, 1999; Mabbott *et al.*, 2013).

M cells are phagocytic epithelial cells that enable the uptake and transcytosis of luminal antigens into the GALT, they are responsible for the rapid transport of bacterial antigens to antigen-presenting immature dendritic cells (Neutra, Frey and Kraehenbuhl, 1996; Rios *et al.*, 2016). M cells display characteristically different morphology from neighboring epithelial cells. On their apical side they have irregular, short microvilli, while on their basolateral side they have a M shaped pocket structure that houses antigen-presenting cells such as macrophages, B cells and dendritic cells (Reynolds, 1987; Vĕtvicka, 1987; Kelsall and Strober, 1996; Iwasaki and Kelsall, 1999). Mature M cells express the receptor Glycoprotein 2 (Gp2), these receptors are critical for the uptake of *Salmonella Typhimurium* and several other pathogenic antigens (Hase *et al.*, 2009). Mice lacking M cells or Gp2 receptors exhibit profound deficiencies in immune response and a decline in the production of antigen specific secretory IgA (s-IgA) in the gut. For instance, absence of M cells or Gp2 expression impaired antigen-specific T cell responses in mice infected with *Salmonella Typhimurium* (Hase *et al.*, 2009; Rios *et al.*, 2016; Kishikawa *et al.*, 2017).

M cells arise from cycling intestinal stem cells in the crypts (which express Lgr5) but they are predominantly localized in the FAE due to the stimulation by nuclear factor κ B ligand (RankL)(Knoop *et al.*, 2009; de Lau *et al.*, 2012). RankL is secreted from specialized stromal cells under the FAE which are known as M cell inducer cells (Nagashima *et al.*, 2017). RankL binds to Rank receptors on the Lgr5+ cells to activate TRAF6 which in turn leads to a signaling cascade of NFκB signaling, both classical and non-canonical (Kanaya *et al.*, 2018). Spi-B and Sox8 were identified as important transcription factors necessary for differentiation and functionality of M cells; Spi-B null and Sox8 null mice showed lack of M cells and severely impaired transcytosis capabilities (Kanaya *et al.*, 2012; Kimura *et al.*, 2019). However, neither Spi-B nor Sox8 were sufficient for Gp2 expression as seen in both Spi-B null mice with intact Sox8 activation and Sox8 null mice with intact Spi-B activation. Although, the progenitor cells approaching the FAE are constantly stimulated by activating signals, only ~10–20% of the cells in FAE are M cells. This perhaps suggests the presence of a regulatory mechanism in the FAE that prevents all the cells from differentiating into M cells in the FAE. Osteoprotegerin (OPG) is a soluble decoy receptor for RankL and it plays a regulatory role in maintaining the M cell density in the intestine by competing with Rank for binding to RankL. OPG null mice exhibited an increase in functionally mature M cell (Kimura *et al.*, 2020). Despite the important role M cells play in initiating mucosal responses, the mechanism for M cell development have yet to be fully characterized.

To further understand the differentiation and development of M cells, we had previously performed Chip-seq and Gro-seq for M cells and discovered 12 transcription factors that were epigenetically regulated (6 upregulated and 6 silenced) (George *et al.*, 2020). One of the genes upregulated in our analysis was Atoh8. We find that Atoh8 plays a critical role in regulating the differentiation of M cells. We found that Atoh8 was expressed exclusively in M cells in the Peyer’s patches and was critical to maintain the density of M cells in the FAE. Intestinal specific Atoh8 deletion showed an increased capacity for transcytosis, suggesting an increased humoral response. However, previous studies have shown that mice with an increase in M cell population have impaired mucosal immunity due to the high volume of bacterial translocation via M cells. Our findings show that Atoh8 plays a homeostatic role in maintaining M cell population in the context of translocation of invasive antigens as well as maintaining the required immune responses from the mucosa.

## Results

### Atoh8 is expressed in M cells and induced by RankL-Rank signaling

In our previous study we sought to identify how PRC2 regulates M cell differentiation and discovered 12 transcription factors that were PRC2-regulated specifically in M cell differentiation. (6 upregulated and 6 silenced) (George *et al.*, 2020). Atoh8 turned up as one of the Rankl-induced PRC2 regulated gene (log2 fold changes −3.15 Rankl vs WENRC and −2.58 RankL vs ENRI) (Fig.1A). Immunohistochemistry analysis of murine PP’s revealed that Atoh8 is localized in the nuclei of cells in the Peyer’s Patches (Fig.1B). RNA was isolated from the FAE and villous epithelium, and RT-qPCR analysis confirmed that Atoh8 was significantly enriched in the Peyer’s patches (Gp2 as marker) when compared to the villi (Fig.1C). Mouse intestinal organoids isolated from the crypts were treated with RankL for 4 days, Atoh8 was observed to be significantly upregulated along with GP2, confirming that Atoh8 is induced by RankL signaling (Fig.1D). Rank-deficient mouse intestinal organoids were generated using Lenti V2 Crispr/Cas9 system, RankL treatment did not induce Atoh8 expression suggesting that Atoh8 expression falls under the purview of Rank-RankL signaling (Fig.1E). Validation of RANK KO was confirmed via immunoblot analysis (supplementary Fig 1 A). Previously in osteoclast, it was shown that BMP induced Atoh8 regulated the RankL/OPG ratio indirectly via Runx2 to regulate osteoclast number and maintain bone volume in mice (Yahiro *et al.*, 2020). Our Gro-seq analysis of RankL organoids from our previous paper revealed that Rankl signaling did lead to an upregulation of BMP2 and BMP6 but not Runx2 (George *et al.*, 2020). The BMP6 and BMP2 activation by Rankl was confirmed by qPCR analysis (supplementary Fig.3A). Our preliminary experiment growing Intestinal organoids in EGF, Rspondin and 100ng/ml of BMP2 and BMP6 according to Calpe et al’s protocol (Calpe *et al.*, 2015) showed that Atoh8 could be induced by BMP2 and BMP6 alone (supplementary Figure 3 C and D).

**Figure 1.**
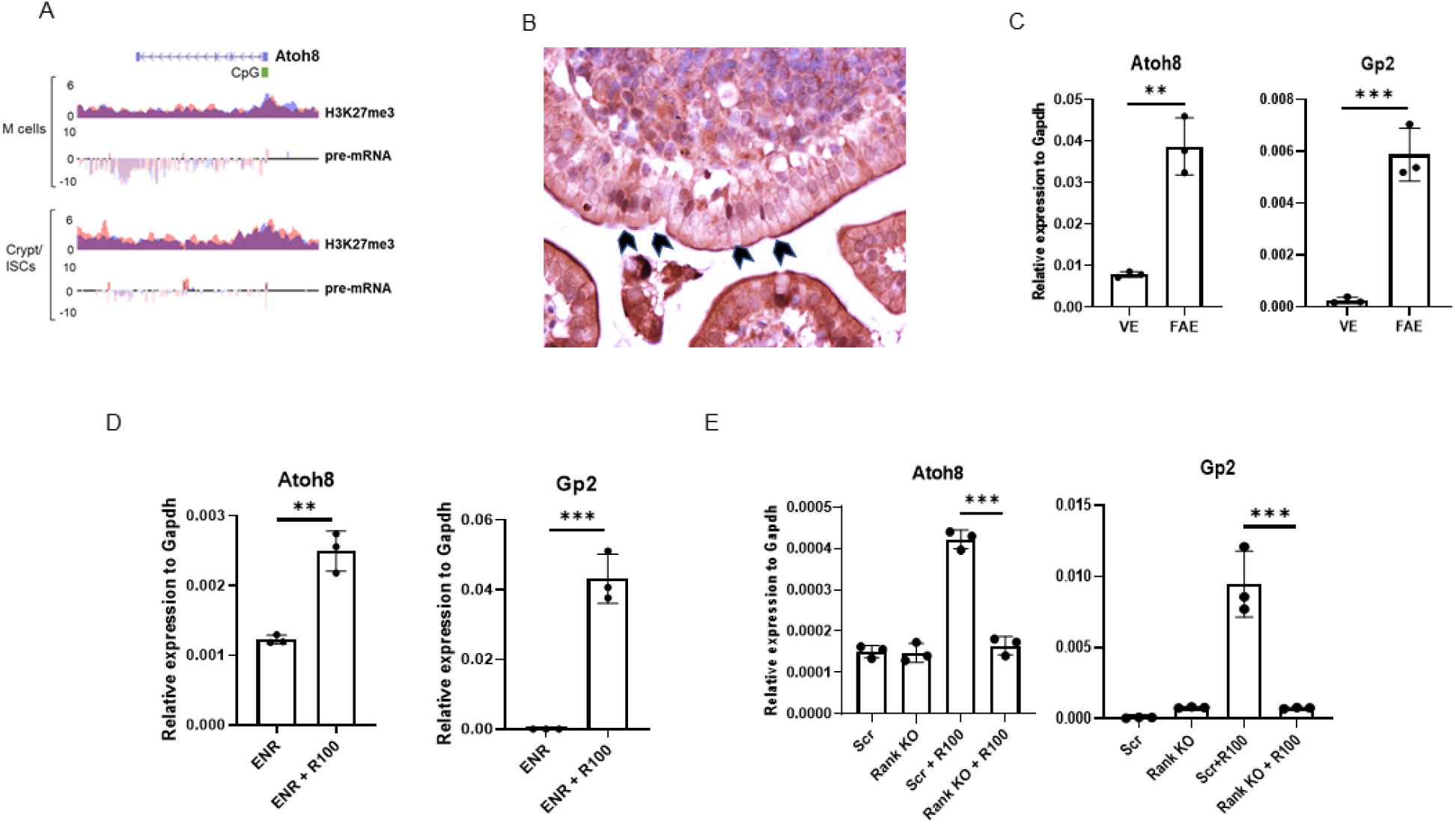
Atoh8 is expressed in FAE in Peyer’s patches and is dependent on Rank-RankL signalling. A) H3K27me3 occupancy at CpG islands spanning the promoter and first exon of the Atoh8 gene in organoids treated with RankL (M cells) or inhibited with IWP2 (Enterocytes) or treated with Wnt3a and Chir99021 (Crypt/ISCs). Below, pre-mRNA expression of Atoh8 in organoids treated as above (y-axis: normalized tag count, ENR500 = R-spondin 500ng/ml, R100 = Rankl 100ng/ml). B) Section of PP from wild-type mice stained with Atoh8 antibody. Arrowheads indicating Atoh8 expression in the nuclei of M cells in FAE. C) RT-qPCR analysis of Atoh8 and Gp2 in the FAE and VE from C57BL/6JRj mice (n = 3). D) Organoids generated from wild-type mice were stimulated with 100ng of RankL for 4 d. Atoh8 and Gp2 expression was examined by RT-qPCR analysis. E) Rank KO organoids and Scrambled organoids generated by lentiCRISPR v2 were incubated with RankL for 4 days, Atoh8 and GP2 expression was analyzed by RT-qPCR. In (C-E) unpaired two-tailed Student’s t test was performed for three independent experiments, ****, P < 0.0005; ***, P < 0.005; **, P < 0.01

### Atoh8 deficiency augments M cell differentiation along with other M cell-associated transcription factors

Since our Chip-Seq and Gro-seq analysis demonstrated that Atoh8 is a PRC2-regulated gene in M cell differentiation, located in PPs and induced by RankL, we hypothesized that Atoh8 may contribute to M cell differentiation. We generated intestine specific Atoh8 knockout by crossing Atoh8 lox/lox with VilCre mice (from now referred as Atoh8 lox/VilCre). M cells were analyzed from the GALT of Atoh8 *lox/VilCre* mice. RT-qPCR analysis of RNA isolated from the FAE of Atoh8 *lox/VilCre* showed an increase in the expression of mature GP2+ M cells, when compared to the control *VilCre* mice (Fig. 2A). Early expression markers of M cells such as CCL20, MarcksL1 and TNFAIP2 were also upregulated (Fig. 2A). Esrrg which is previously known to be essential for development of M cells was also shown to be upregulated (Fig.2A). Spi-B and Sox8, which are transcriptional factors critical for functional M cell development, were also markedly higher in the Atoh8 *lox/VilCre* mice (Fig. 2A). OPG, the decoy receptor for RankL was also showed higher expression in the Atoh8 lox/VilCre mice. The increase in Gp2+ M cells was also corroborated with whole mount immunostaining (Fig. 2B). We observed increased Gp2+ cells in the Atoh8 *lox/VilCre* mice, counting 22.4 ± 2.85 cells/0.01mm in comparison with control Vil/Cre mice that had 15± 0.09 cells/0.01mm

**Figure 2:**
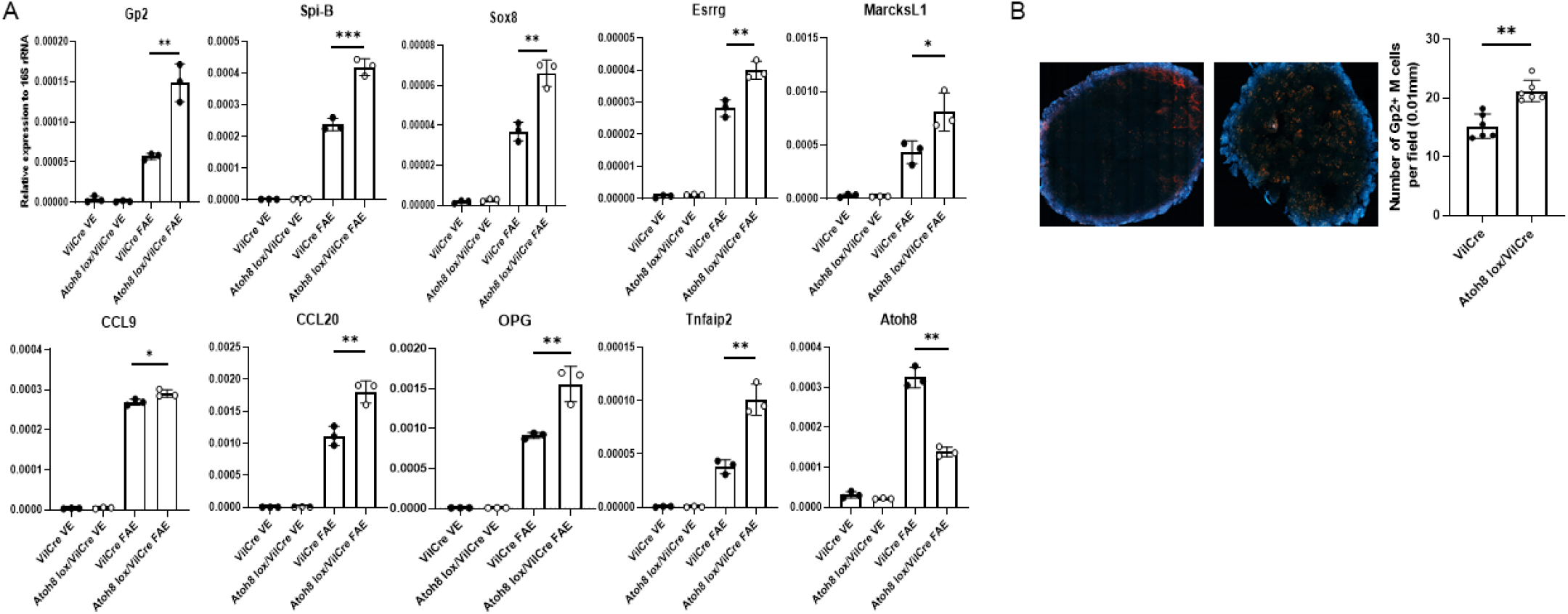
The loss of Atoh8 increases the number of mature M cells in the FAE. (A) qPCR analysis of M cell–associated genes in the FAE and VE of Atoh8 lox/VilCre and VilCre mice. **, P < 0.01; n.s., not significant; unpaired two-tailed Student’s t test, n = 3. (B) Whole-mount immunostaining of PPs from Atoh8+/− and Atoh8−/− mice. The number of GP2+ M cells per field of FAE (0.01 mm2) was compared between them (n = 8). Scale bars, 200 μm.

### Atoh8 deficiency did not affect the composition of lymphoid tissues in the FAE

Germinal centers (GC) in the GALT undergo multiple changes in terms of somatic hypermutation and class switch recombination, this leads to a constant change in the population of lymphoid cells (Muramatsu *et al.*, 2000; Kawamoto *et al.*, 2012). Knockdown of transcription factors necessary for M cell differentiation like Spi-b and Sox8 lead to lack of Gp2 cells as well as reduced B and T cell population in the Peyer’s patch (Kanaya *et al.*, 2012; Kimura *et al.*, 2019). OPG is another gene upregulated in M cell development. The gene transcribes a soluble decoy receptor that binds to RankL. Knockdown of OPG exhibited an increase in M cell population as well as an increase in lymphoid cells that are involved in mucosal response(Kimura *et al.*, 2020). The increase in functionally mature M cells in Atoh8 *lox /VilCre* mice would indicate that the mice would perhaps have an increase in systemic lymphoid tissues and enhanced immune response. Strikingly we did not observe any increased production of high affinity IgA+ in the Atoh8 *lox/VilCre* mice. Similar population of total B cells and T cells were observed in both sets of mice at 4 weeks of age (Fig. 3.A-C). We also looked at follicular T helper cells (Tfh cells) since they promote germinal center reactions and affinity maturations to produce high affinity IgA (Tsuji *et al.*, 2009). Tfh cells and GC B cells had similar population in *VilCre* and Atoh8 *lox/VilCre* (Fig.3C). Since there wasn’t much change in the GC, no differences were observed in the IgA class switched B cells either. RankL producing Th cells retained similar population numbers in both sets of mice.

**Figure 3.**
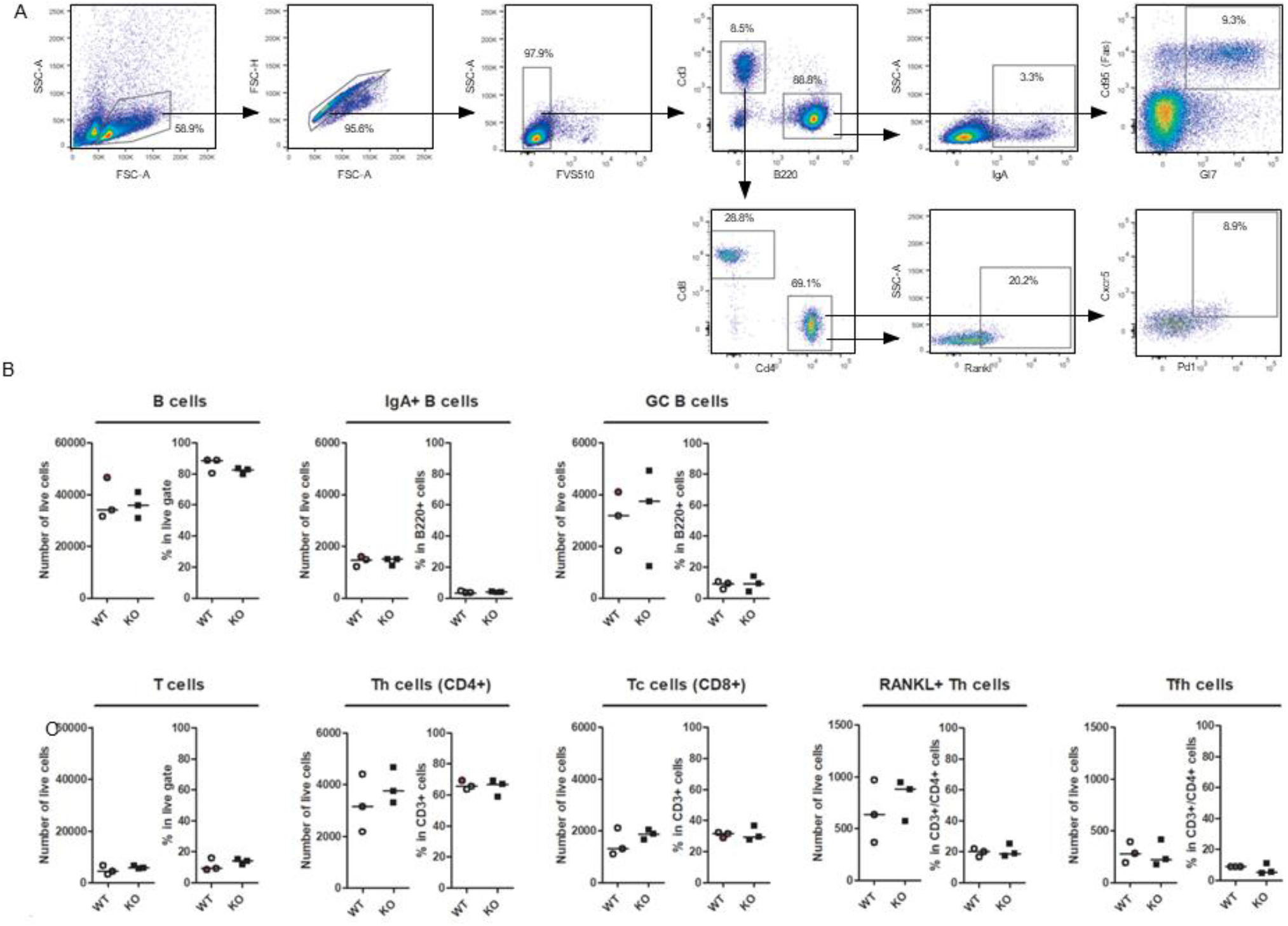
The loss of Atoh8 did not change the composition of lymphoid cells in 4-week-old-old mice after weaning. (A) Gating scheme for analysis of B and T cell populations in VilCre and Atoh8 lox/VilCre ileal PPs. B (CD3ε−B220+) cells were analyzed for IgA+ B and CD95+GL7+ GC B cells. T (CD3ε+B220−) cells were analyzed for total CD4+ Th, CD8α+ cytotoxic T (Tc), and CD4+CD8α−CXCR5+PD-1+ Tfh cells. (B and C) FlowJo analysis of indicated immune cells in ileal PPs. (B) Number of total B cells, GC B cells, and IgA+ B cells. (C) Number of total T cells, Tc cells, Th cells, and Tfh cells; n.s., not significant; Student’s t test, n = 3 per group.

### Epithelium intrinsic Atoh8 is responsible for increase in M-cell population

The subepithelial dome is an afront to multiple signaling pathways, therefore we thought it was necessary to validate if Atoh8 regulation of M cell development was epithelium intrinsic. We established a mouse intestinal organoid culture isolated from crypts of Atoh8 *lox/VilCre* and *control VilCre* mice. These organoids were treated for 4 days with 100ng/ml of RankL to induce M cell differentiation. Spi-B, Sox8 and Esrrg were observed to be upregulated significantly from the control. The epithelial cells were able to mimic the significantly increased Gp2 expression as observed in vivo in the FAE as well as the early markers expressions of CCL9, Marcksl1 and TNFAIP2 (Fig.4A). Immunostaining images for Gp2 on intestinal organoids treated with RankL also showed higher expression for Atoh8 lox/VilCre organoids compared to *VilCre* organoids (Fig.4B). Taken together, these data indicate that epithelial intrinsic Atoh8 is sufficient to regulate M cell differentiation.

**Figure 4.**
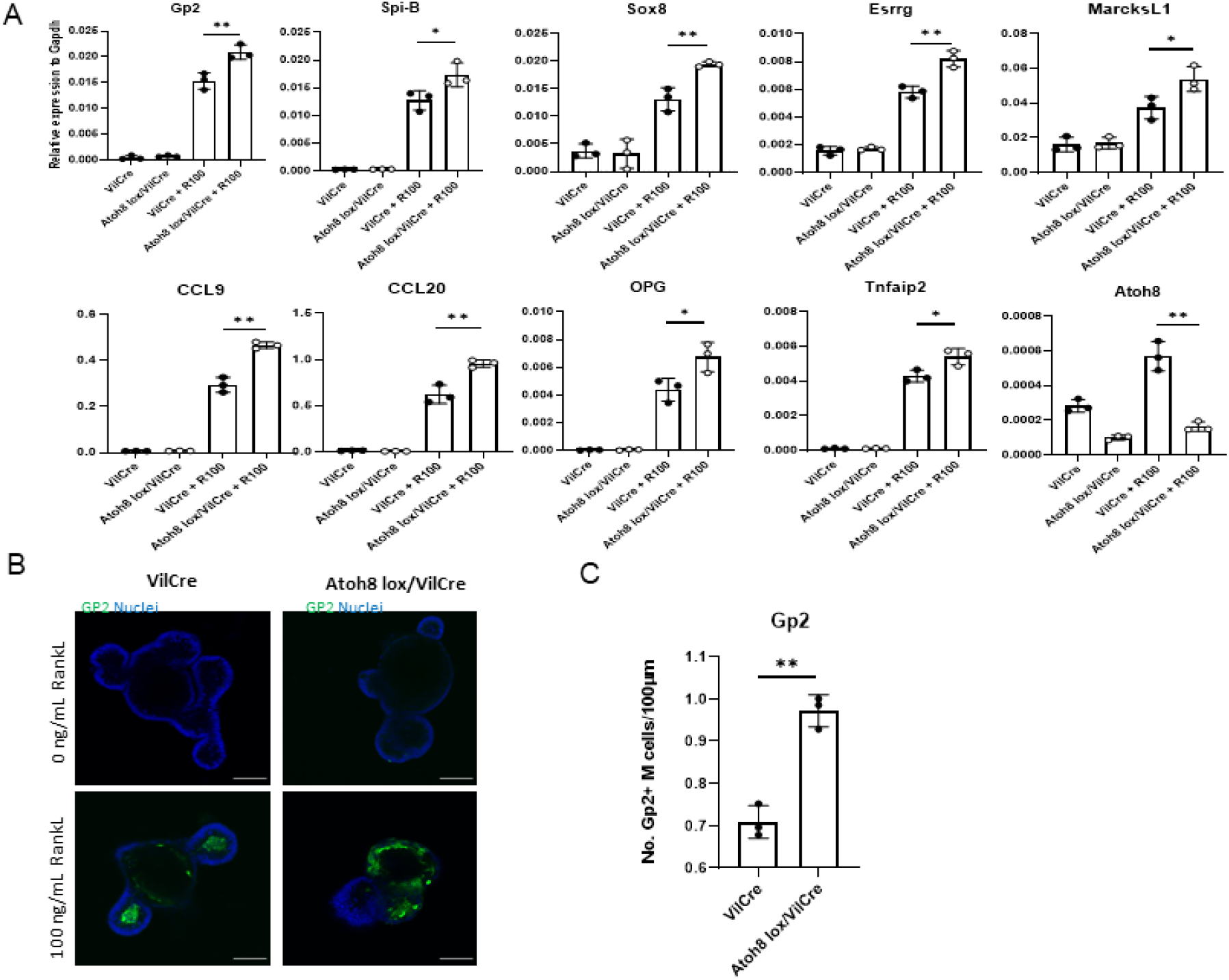
Increase in M cell maturation in Atoh8 lox/VilCre results from epithelium-intrinsic defect. (A and B) Organoids from small intestinal crypts of Atoh8 lox/VilCre and VilCre mice were cultured with or without RANKL for 3 d. (A) qPCR analysis of M cell–associated genes expressed in the organoid cultures. Values are presented as the mean ± SD; **, P < 0.01; *, P < 0.05; n.s., not significant; unpaired two-tailed Student’s t test, n = 3. Data are representative of two independent experiments. (B) Immunostaining images for GP2 (green) on organoids. Bars, 100 μm. E) The number of Gp2+ M cells and Sox8+ M cells per length epithelium of organoids was compared between VilCre and Atoh8 lox/VilCre treated organoids (n = 3). Images are representative of three independent experiments.

### Atoh8 deficiency leads to increased transcytosis capacity

Gp2+ M cells in the Peyer’s patches are essential for transcytosis of antigens and foreign particles to initiate mucosal immune response. Since Atoh8 knockout mice presented more Gp2 cells in the FAE, we investigated if these Gp2 cells are functional. To explore this, we administered orally fluorescent nanoparticles to Atoh8 *lox/Vil-Cre* and control *Vil-Cre* mice. 4 hours after the oral administration, PP’s from both mice were removed and the number of particles transcytosed in each PP was counted with a fluorescence microscope (Fig.5A). We observed that the uptake of nanoparticles in the Atoh8 *lox/VilCre* Peyer’s patch was increased significantly by over 2-fold when compared to the *control* mice (Fig.5B).

**Figure 5.**
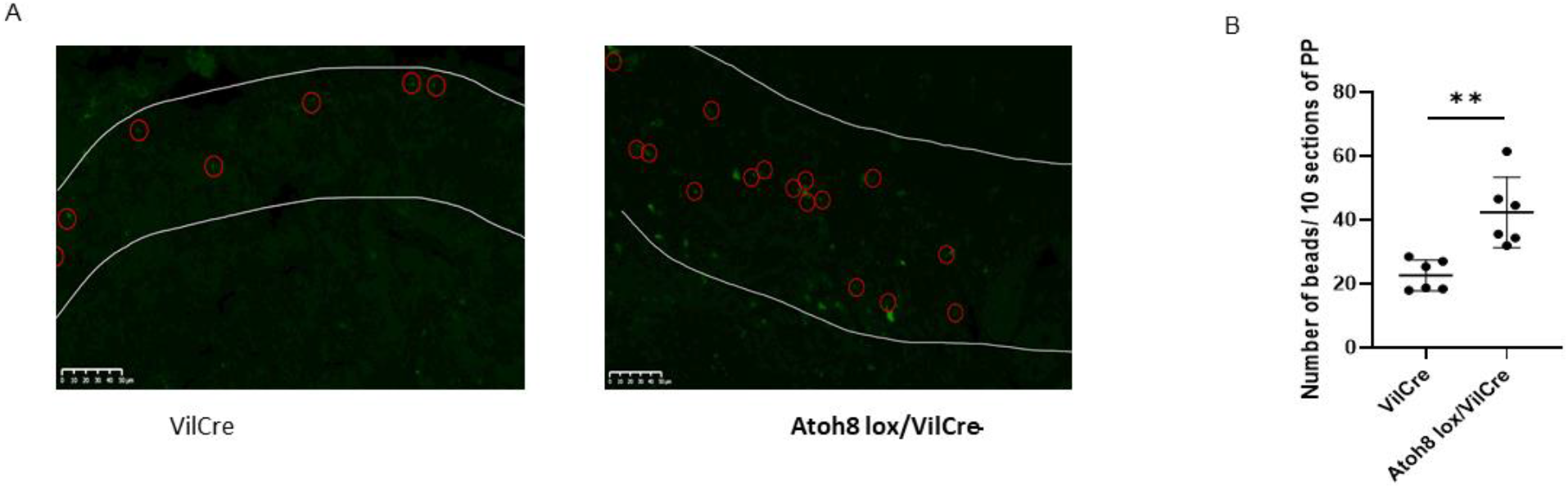
The loss of Atoh8 causes increased uptake of luminal nanoparticles into the follicle. (A and B) Green fluorescent latex beads (200-nm diameter) were orally administered to Atoh8+/+ or Atoh8−/− mice. 3 h later, PPs were collected from the jejunum. Ten consecutive cryosections of each PPs were examined by fluorescence microscopy, and the number of particles was counted manually. (A) Representative images of cryosections. White lines indicate FAE. Red circles indicate fluorescent particles. Bars, 50μm. (B) Quantification of particles in PPs. **, P < 0.01; Student’s t test; n = 3 per genotype.

## Discussion

M cells are Intestinal epithelial cells specialized in gut immunity and transcytosis of antigens via GP2 to initiate immune responses. Previous research has revealed that differentiation into M cell requires activation of Nf-κB signaling, activation of transcription factors Spi-B, Esrrg and Sox8, membrane bound RankL from lymphoid cells and S100A4 from Dock8 cells (Kanaya *et al.*, 2012, 2018; Kimura *et al.*, 2019; Kunimura *et al.*, 2019). However, other factors controlling M cell developments remains to be elucidated. For our analysis into M cell differentiation, we had previously performed a Chip-seq and Gro-seq of M cells and our data set revealed that PRC2 regulated 12 novel transcription factors (6 silenced and 6 upregulated) in M cell development (George *et al.*, 2020). Atoh8 was one of the 6 transcription factors regulated by the PRC2. From our study we describe that M cells in the FAE expressed Atoh8, it was also specifically induced by RankL treatment in intestinal organoids differentiating to M cells and came under the purview of RankL-Rank signaling pathway.

Atoh8 belongs to the group A of the bHLH transcription factor family. Members of the Atonal superfamily control numerous aspects of differentiation for vertebrae organ development and function (Yan, 1998; Tomita *et al.*, 2000; Hutcheson and Vetter, 2001). Atoh1 is a member of the bHLH family that plays a pivotal role in differentiation of Paneth cells. Paneth cells are critical for providing stem cell niche to Lgr5+ cells in the intestinal crypts (Durand *et al.*, 2012). Atoh8 is the sole mammalian member of the bHLH factor that is a part of the NET family (Rawnsley *et al.*, 2013). Atoh8 has been previously studied in the context of differentiation of osteoblasts. In a recent study Atoh8 was found to be induced by BMP signaling. BMP induced Atoh8 regulated the RankL/OPG ratio indirectly via Runx2 to regulate osteoclast number and maintain bone volume in mice (Yahiro *et al.*, 2020). In our preliminary experiment with RankL treated intestinal organoids we observed that RankL did give rise to BMP2 and BMP6 expression. However, Runx2 was not activated by RankL. We were also able to observe that Atoh8 could be induced by BMP2 and BMP6 alone. However, since Runx2 was not activated by RankL, further investigation is required to find out if BMP2/BMP6 directly induces Atoh8 to control the density of M cell population in the Peyer’s patch.

Our Atoh8 intestinal specific knockout mouse exhibited a higher number of Gp2+ M cells when compared to the VilCre wildtype. The well-established early markers of M cells such as MarcksL1, TNFAIP2 were all significantly upregulated. Spi-B and Sox8, which are critical for the maturation of Gp2+ M cells and fall under the purview of RankL and Nfkb (both classical and non-canonical) signaling were found to have higher transcriptional expression. Isolation of intestinal organoids and treatment with RankL indicated that Atoh8 deficiency that led to increased Gp2 was epithelium intrinsic. Previous research involving transcription factors involved in M cell differentiation like Spi-B and Sox8 showed that transcription factor deficient mice also exhibit reduced lymphoid cell population in PP along with reduced GP2 expression (Kanaya *et al.*, 2012; Kimura *et al.*, 2019). Osteoprotegrin (OPG) deficient mice showed higher Gp2+ M cells but also higher numbers for lymphoid cells in terms of population (Kimura *et al.*, 2020). The regulation of M cell density by Atoh8 did not affect antigen-specific mucosal and systemic lymphoid population mediated by IgA. The population of lymphoid cells in the Peyers patch of Atoh8 lox/VilCre remained unchanged after weaning despite the increased Gp2+ M cell. To account for delayed immune response/population changes, 10-week-old Atoh8 lox/VilCre mice didn’t show any changes in lymphoid population either when compared to the control (Supplementary Fig.4). However, further studies with Salmonella Typhimurium infection or other pathogenic antigen could induce an enhanced immune response. Atoh8 intestinal KO exhibited a higher transcytosis capacity of nanobeads into the Peyer’s patches, twice the amount of uptake when compared to control mice. This indicates that Atoh8 lox/VilCre mice with higher Gp2 expression would transcytosis higher numbers of antigen during an infection.

OPG binds to RankL and acts as a decoy receptor instead of Rank. OPG deficient mice showed higher RankL secretion, more Gp2+ M cells, higher transcytosis capabilities and increased population of systemic lymphoid tissues and enhanced immune response (Kimura *et al.*, 2020). We looked to see if Atoh8 intestinal KO mice had increased RankL production, our FACS analysis showed no increase in RankL+ T cells, indicating that Atoh8 acts independent of RankL signaling in the GALT. Although the OPG null mice showed higher M cells and enhanced immune responses, they were highly susceptible to infection by pathogenic bacteria. An increase in M cells could also increase the transcytosis of botulinum toxins and scrape prion protein into the body. (Matsumura *et al.*, 2015) Therefore, considering that the lack of Atoh8 in the intestine led to an increase in M cells, it is reasonable to reason that Atoh8 limits transcytosis of pathogenic agents by regulating the number of M cells. The delicate equilibrium of maintaining M cell density in the Peyer’s patch via Atoh8 may have been established over time as an evolutionary mechanism to control intestinal homeostasis in relationship to invasive antigens and mucosal immune responses. Atoh1, a family member of Atoh8 regulates Paneth cell differentiation in the intestinal crypt via notch signaling and simultaneously inhibiting Paneth cell differentiation in the neighboring cell perhaps Atoh8 employs a similar mechanism in maintaining the population. However, further exploratory studies are needed to understand how the epithelial intrinsic Atoh8 regulates the population of M cells.

## Materials and Methods

### Mice

All animal experiments were approved by the Finnish National Animal Experiment Board (Permit: ESAVI/5824/2018). Mice were maintained on standard light-dark conditions, with food and water ad libitum at the pathogen free animal facility of the faculty of Medicine and Health Technology. B6.Cg-Tg(Vil1-cre)1000Gum/J mice (Cat No: 021504) were purchased from Jackson Laboratories. Atoh8 lox/lox mice, in which exon 1 is flanked by two loxP sites, were provided by Rosa Gasa (Ejarque, et al. 2016). To generate intestinal deletion of Atoh8, Vil1-cre mice were bred with Atoh8 lox/lox. F1 generation was backcrossed with Atoh8 lox/lox to generate mice homozygous for floxed Atoh8 allele carrying the Vil1-cre. Littermates with Vil1-cre allele were used as control. Mice genotypes and Atoh8 deletion was confirmed by PCR.

### Immunohistochemistry and immunofluorescence

Peyer’s patches were isolated from the ileum and washed in cold PBS and embedded into paraffin blocks. Sections were cut from the blocks and rehydrated by washing with PBS. After blocking with 1% PBS/BSA supplemented with 5% normal donkey serum, antigen retrieval was processed with citrate buffer, pH 6.0 (121°C for 5 min) and stained overnight at 4°C for Atoh8 (Thermo, ab49129) antibody. Anti-Rabbit was used for the secondary antibody. Light microscopy was used for detection and analysis.

Intestinal crypt organoids were analyzed by whole-mount immunostaining, crypt-organoids were grown in an 8-well chamber plate and cultured for 4 days with and without RankL 100ng/ml after which they were fixed with 4% PFA, followed by permeabilization with 0.1% Triton X-100. The organoids were stained with Gp2 (MBL, D278-3) antibodies overnight at 4 degrees Celsius. This was followed by Anti-Rat for the secondary antibody. Gp2 expressing cells were analyzed by Nikon A1R+ Laser Scanning Confocal Microscope after mounting with ProLong Diamond with DAPI mounting solution (Molecular Probes P36962).

### Isolation of follicle associated epithelial cells (FAE) and villous epithelium cells (VE)

Illeal PP’s along with small pieces of intestine were isolated from the ileum of control mice and Atoh8 lox/VilCre mice. After flushing the tissues with cold PBS, they were incubated in 30mM EDTA, 5mM DTT in PBS and gently shaken in ice on a rocker for 20 minutes. Surrounding epithelial cells were peeled off from lamina propria and PP’s. FAE was carefully cleaned off from surrounding VE tissues with a 26-gauge needle under a stereo microscope. Trizol was added to the cleaned FAE and proceeded by RNA isolation.

### Mouse Intestinal Organoid Culture

Mouse intestinal crypts were isolated and culture techniques were observed as previously described by Sato et al 2011 and de Lau et al 2012 17,29. Collected duodenum were washed in PBS and cut longitudinally, villi were gently scraped off using glass slides. After further washes with PBS, the duodenum was cut into 2mm pieces, pipetted up and down 4-6 times in 10 ml PBS using a 10ml pipette. Once the suspension was relatively clear, the pieces were suspended in in 10mM EDTA in PBS for 20 minutes rocking at room temperature. A 70-μm cell strainer (Fisher Scientific) was used to strain the crypts from the rest of the epithelium. This mixture was enriched to crypt fraction through centrifugation at 150 × g for 5 minutes. The crypts were cultured on a 24 well plate by embedding them in 30ul of Matrigel (Corning). Organoids were cultured in an optimal medium consisting of advanced DMEM/F12 (Thermo Fisher Scientific) that contained HEPES (10mM, Sigma-Aldrich), Glutamax (2mM, Thermo Fisher Scientific), Penicillin-streptomycin (100U/ml, Sigma-Aldrich), B-27 supplement minus Vitamin A (Thermo Fisher Scientific), N-2 supplement (Thermo Fisher Scientific), N-acetylcysteine (1 mM; Sigma-Aldrich), recombinant WNT (100ng/ml R&D biosystems), recombinant murine EGF (50 ng/mL; Thermo Fisher Scientific), recombinant murine Noggin (50 ng/mL; PeproTech), recombinant mouse R-spondin1 (1 μg/mL; R&D Systems). Media were changed every 2 days. For M cell differentiation, recombinant mouse RankL (100ng/ml, Peprotech) was added to the media and incubated for 4 days. To check for Atoh8 expression via BMP2/BMP6, organoids were grown in EGF, Rspondin and BMP2 (100ng/ml, R&D biosystems) and EGF, Rspondin and BMP6 (100ng/ml, R&D biosystems). Noggin was removed for the BMP2/BMP6 experiments as it acts as an antagonist against BMP signaling.

### CRISPR–Cas9 gene knockout of intestinal organoids

Guide RNA’s for the Rank gene were designed using CRISPR design tool (http://crispr.mit.edu). (Shalem, O. et al. Science 343, 84–87 (2014)). The guides were cloned into lentiCRISPR v2 vector (Addgene, 52961). The cloned product was transfected into HEK 293FT cells (ThermoFisher R7007). After 48 hours the supernatant was collected and concentrated with Lenti-X concentrator (Clontech). The 293FT cell line was tested for mycoplasma. Intestinal organoids were grown in ENCY (EGF, Noggin, Chir-99021 and Y-27632) 2 days prior to transduction. After dissociating organoids into single cells using TrypLE Express (Thermo Fisher Scientific) supplemented with 1,000 U/ml DnaseI for 5 min at 32 °C, the cells were washed once with Advanced DMEM and resuspended in transduction medium (ENR media supplemented with 1mM nicotinamide, Y-27632, Chir99021, 8 μg/ml polybrene (Sigma-Aldrich) and mixed with concentrated virus. The mixture was centrifuged for 1 hr at 600 × g 32 °C followed by 3 hr incubation at 37°C, after which they were collected and plated on 60% Matrigel overlaid with enriched transduction medium without polybrene. On day 2 and day 4, transduced organoids were selected with 2 μg/ml of puromycin (Sigma-Aldrich). After which clones were expanded in maintenance ENR medium. Knockout was confirmed by western blot to check for the expression of deleted gene.

Oligonucleotides used for generation of gRNAs for Rank CACCGAAAGCTAGAAGCACACCAG, AAACCTGGTGTGCTTCTAGCTTTC.

### Real-time quantitative reverse transcription PCR

Total RNA was isolated from intestinal organoids, FAE and VE tissues from mice were using TRIzol (Life Technologies). iScript cDNA synthesis Kit (Biorad, 1708891) was used to transcribe the isolated RNA to first strand cDNA. qPCR amplification was detected using Ssofast evergreen supermixes (Biorad,172-5203). The specific primers used, are listed in the supplementary data.

### Whole mount immunostaining of M cells in FAE

Ileal PPs were dissected from the small intestine and transferred to a 10 cm dish containing 30 ml cold PBS. Excess intestinal tissues around the FAE were cut and removed under a stereo microscope using forceps. PPs were washed sufficiently by using 1 ml syringe with 26 gauge needle under a stereo microscope. Mucous layer on the FAE should be flushed out with water stream to prevent background noise detection. PPs were transferred to the 1.5 ml tube containing 1 ml PBS and washed well by vortexing and the supernatant were discarded. After 3 washes, 300-1000 ul Cytofix/Cytoperm buffer (BD Biosciences) was used for blocking and permeabilization for 25 min at room temperature. Perm/wash buffer (BD Biosciences) was used to pipette wash the PP’s after which they were stained with PE-conjugated anti-GP2 antibody (MBL; 1:10 in Perm/Wash buffer) overnight at 4°C. PP’s were washed 3 times with wash/perm buffer and stained for 30 minutes at RT in Alexa Fluor 546-conjugated anti-Rat IgG (Invitrogen). The PP’s were washed again 3 times with wash/perm buffer and mounted with ProLong Diamond with Dapi mounting solution (Molecular Probes P36962). Slides were examined with a laser scanning confocal microscope (Zeiss LSM 800 LSCM).

### Flow cytometry

Peyers patches from Vil mice and Atoh8 loxp/VilCre were isolated from the ileal section. The tissues were washed in PBS and incubated in 5 ml Spleen Dissociation Medium for 15-20 min at 37 °C while vigorously shaking at 250 rpm. To generate a single cell suspension, PPs were placed on a sterile (autoclaved) 70 μm nylon mesh cell strainer and ground into the mesh using the base of a plunger from a 1 cc syringe. The suspension was incubated in 1mm EDTA for 5 mins in a rocker after which the suspension was passed through a 70 μm nylon mesh cell strainer. After the isolation step, the single cell suspensions were prepared with FVS510 viability stain (#564406; Becton, Dickinson and Company, New Jersey, USA) and CD16/CD32 Monoclonal Antibody (#16-0161-85; Thermo Fisher Scientific, Massachusetts, USA). Surface staining was done using fluorochrome conjugated anti-mouse antibodies against CD3e, CD8a, CD4, B220, PD1, CXCR5, GL7, FAS, IgA and RANKL (Thermo Fisher Scientific). In order to prevent spectral overlap of emitted fluorescence, cells were divided into replicates and two separate antibody panels used in the staining procedure. Flow cytometry was done with FACSAria™ Fusion (Becton, Dickinson and Company), and data were analyzed with FlowJo (v. 10.6.1, Tree Star, Oregon, USA). Statistical analyses were done with the Prism v. 5.02 (GraphPad Software, California, USA) calculated using a nonparametric two-tailed Mann-Whitney OR a two-tailed t-test. P values of < 0.05 were considered significant.

### Quantification of transcytosis of fluorescent beads by M cells

Atoh8 lox/VilCre and VilCre mice were fasted for 3 hours, 10^11^ of 200-nm diameter polystyrene nanoparticles (Fluoresbrite YG; Polysciences) were orally administered via oral gavage. After 4hr, two PP’s were collected from the ileum and jejunum and fixed with 3.7% formalin/PBS for 2 h. Fixed tissues were incubated overnight with 30% sucrose/PBS and finally embedded in OCT compound (Sakura Fintech). Ten sequential 15um sections were cut and examined by fluorescence microscopy. The number of fluorescent particles transcytosed was counted manually.

## Abbreviations

M cells: = Microfold cells;
GALT: = Gut associated lymphoid tissues;
PP: = Peyer’s Patch;
FAE: = Follicle associated epithelium;
VE: = villous epithelium;
RankL: = Receptor activator of nuclear factor kappa B ligand;
Rank: = Receptor activator of nuclear factor kappa B;
PRC2: = polycomb repressive complex 2;
Esrrg: = Estrogen-related receptor gamma.

---

**Supplementary Figure 1.**
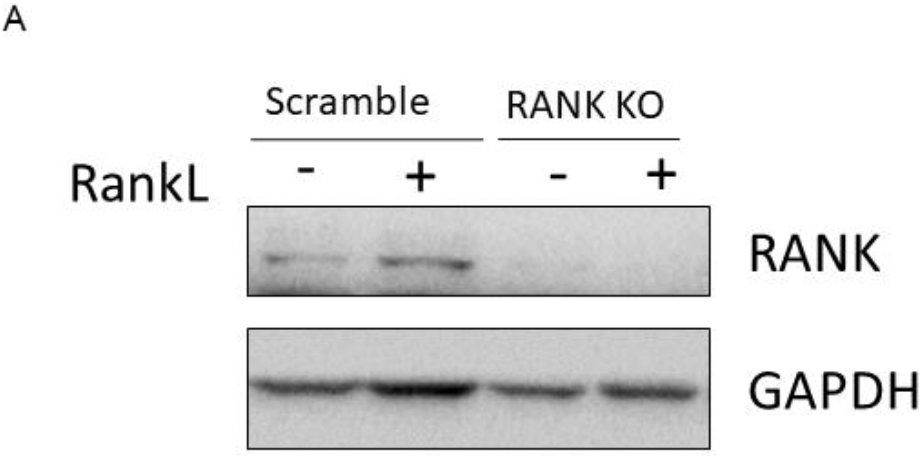
Immunoblot analysis of CRISPR generated RANK KO. A) RANK KO validation by immunoblot analysis of RANK KO and Scrambled intestinal organoids generated by CRISPR-Cas9 genome editing in C57BL/6rj intestinal organoids. These organoids were grown with and without RankL 100ng for 4 days.

**Supplementary Figure 2.**
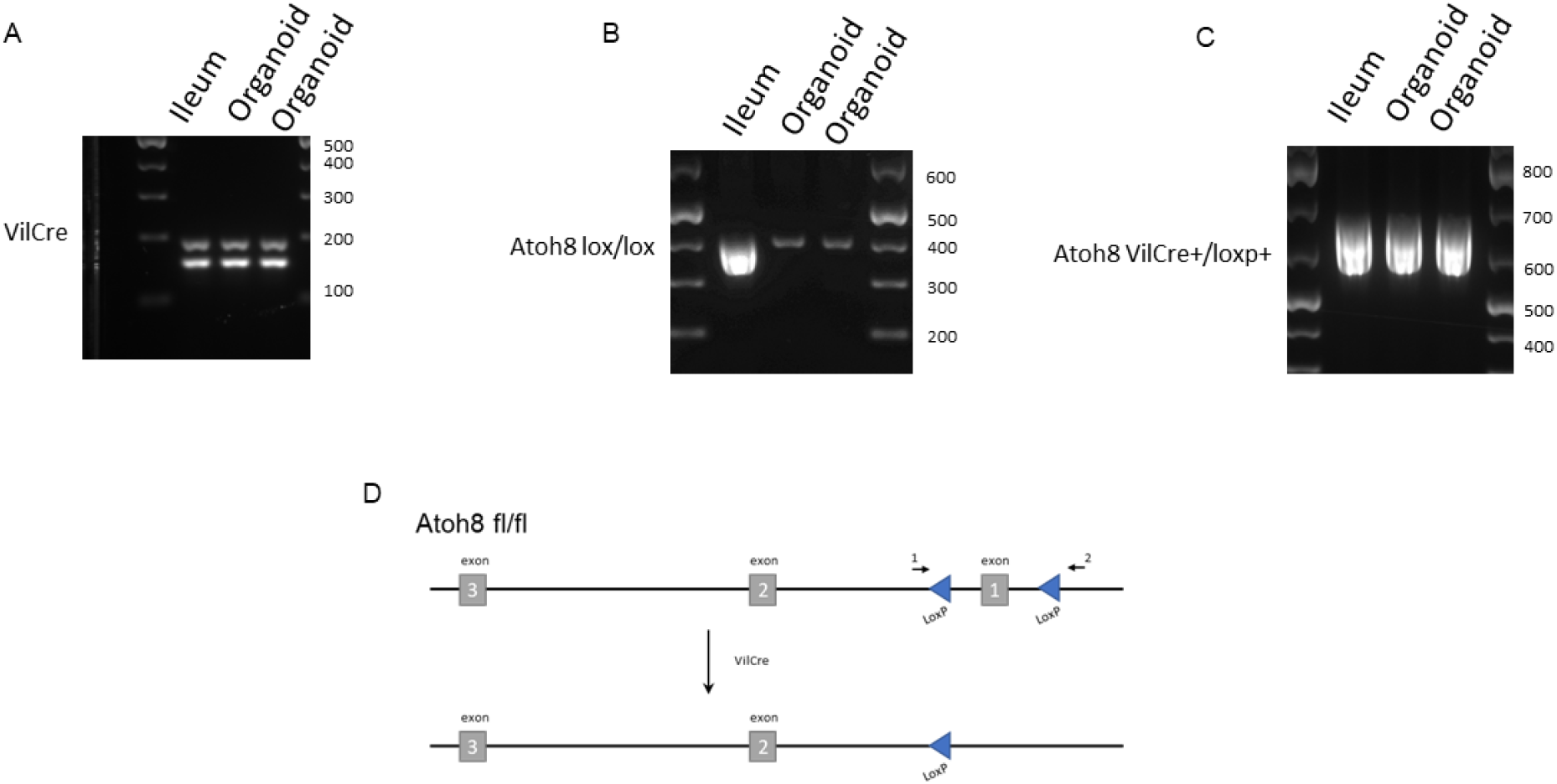
Representative genotyping PCR amplification of intestinal VilCre, Atoh8 flox/flox wild-type and Atoh8 VilCre/loxp. A) PCR for detection of VilCre recombinase is shown. Upper band: internal control (182 bp). Lower band: VilCre transgene (150 bp). 100 bp molecular marker brightest line corresponds to 500 bp. B) Floxed allele 400 bp C) Genotyping of Atoh8 lox/VilCre 600bp D) Scheme and representative PCR amplification verifying Cre-mediated recombination of the Atoh8 floxed allele in the intestine.

**Supplementary Figure 3.**
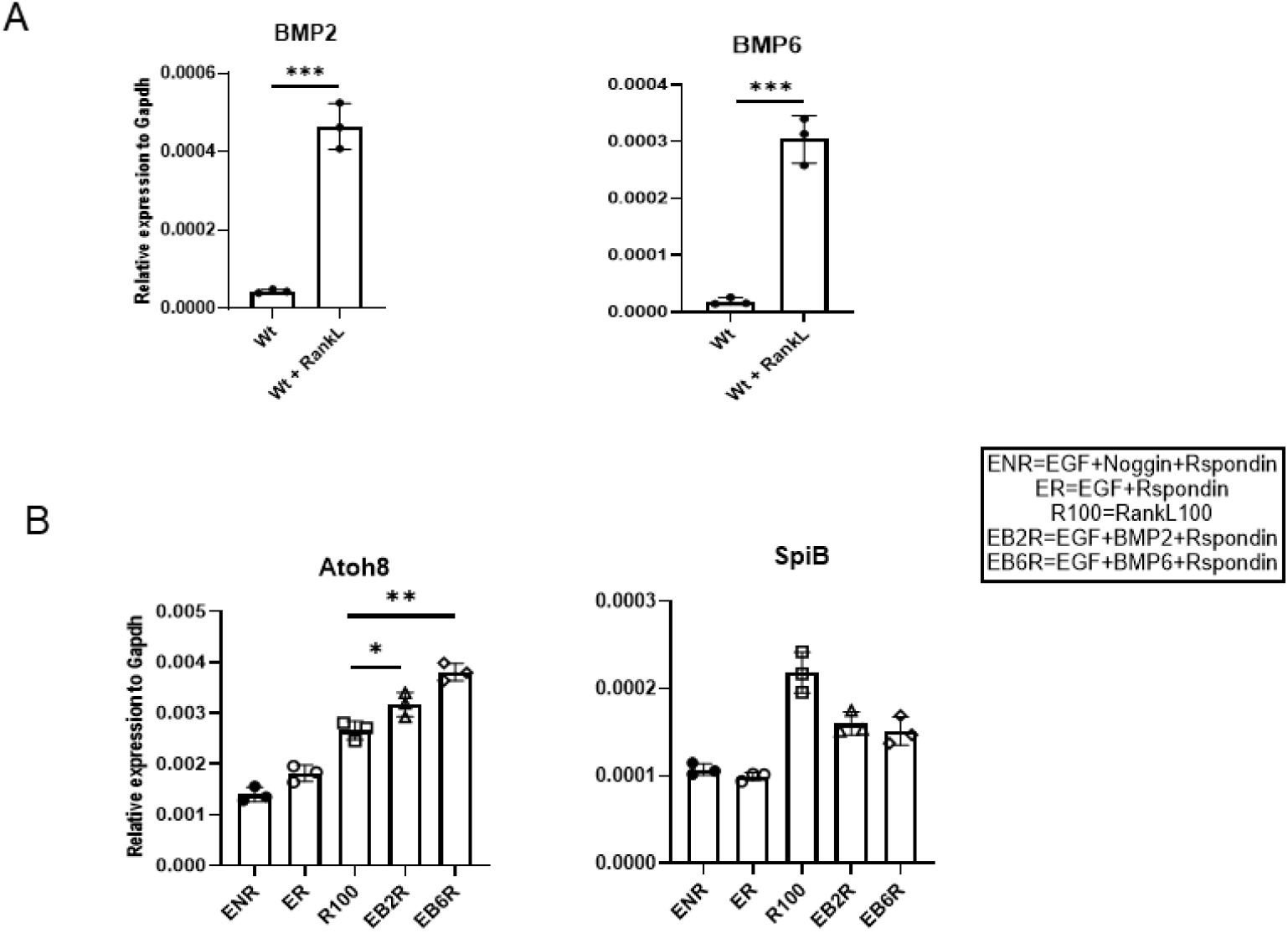
Atoh8 signaling is mediated by RankL/BMP2-BMP6 signaling. (A) Organoids from small intestinal crypts of Wildtype mice were cultured with or without RANKL for 3 d. (A) qPCR analysis of M cell–associated genes expressed in the organoid cultures. Values are presented as the mean ± SD; ***, P < 0.005; unpaired two-tailed Student’s t test, n = 3. Data are representative of two independent experiments. (B) Organoids isolated Wt mice were grown for 3 days in EGF, Noggin, Rspondin media, EGF, Rspondin media, 100ug of Rankl media, Egf, BMP2 Rspondin media and Egf,BMP6, Rspondin media. Values are presented as the mean ± SD; *,P < 0.05, **, P < 0.001; unpaired two-tailed Student’s t test, n = 3. Data are representative of two independent experiments.

**Supplementary Figure 4.**
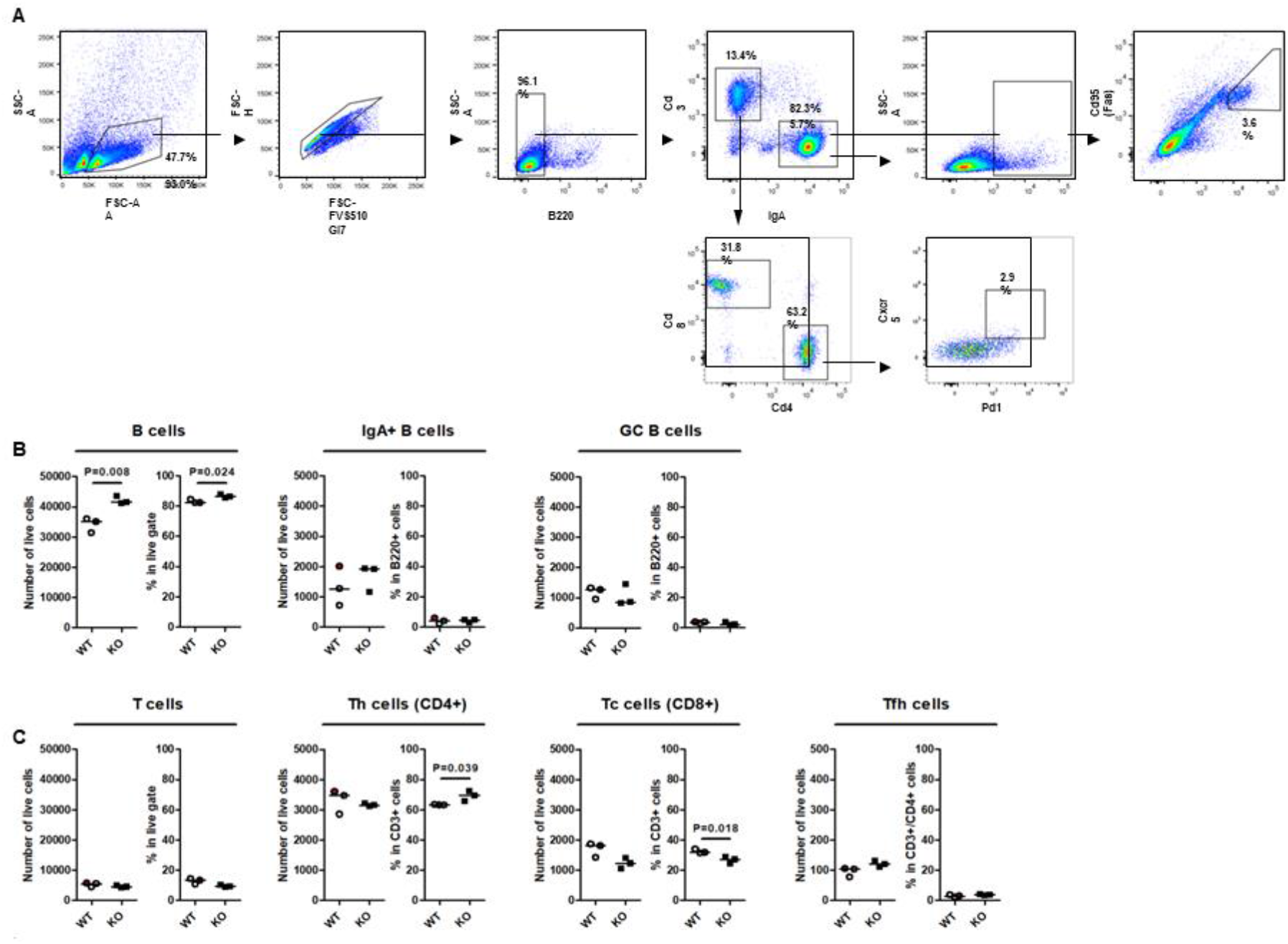
The loss of Atoh8 did not change the composition of lymphoid cells in 10-week-old mice. (A) Gating scheme for analysis of B and T cell populations in VilCre and Atoh8 lox/VilCre ileal PPs. B (CD3ε−B220+) cells were analyzed for IgA+ B and CD95+GL7+ GC B cells. T (CD3ε+B220−) cells were analyzed for total CD4+ Th, CD8α+ cytotoxic T (Tc), and CD4+CD8α−CXCR5+PD-1+ Tfh cells. (B and C) Flow cytometry analysis of indicated immune cells in ileal PPs. (C) Number of total B cells, GC B cells, and IgA+ B cells. (D) Number of total T cells, Tc cells, Th cells, and Tfh cells; n.s., not significant; Student’s t test, n = 3 per group.

**Supplementary Table 1.**
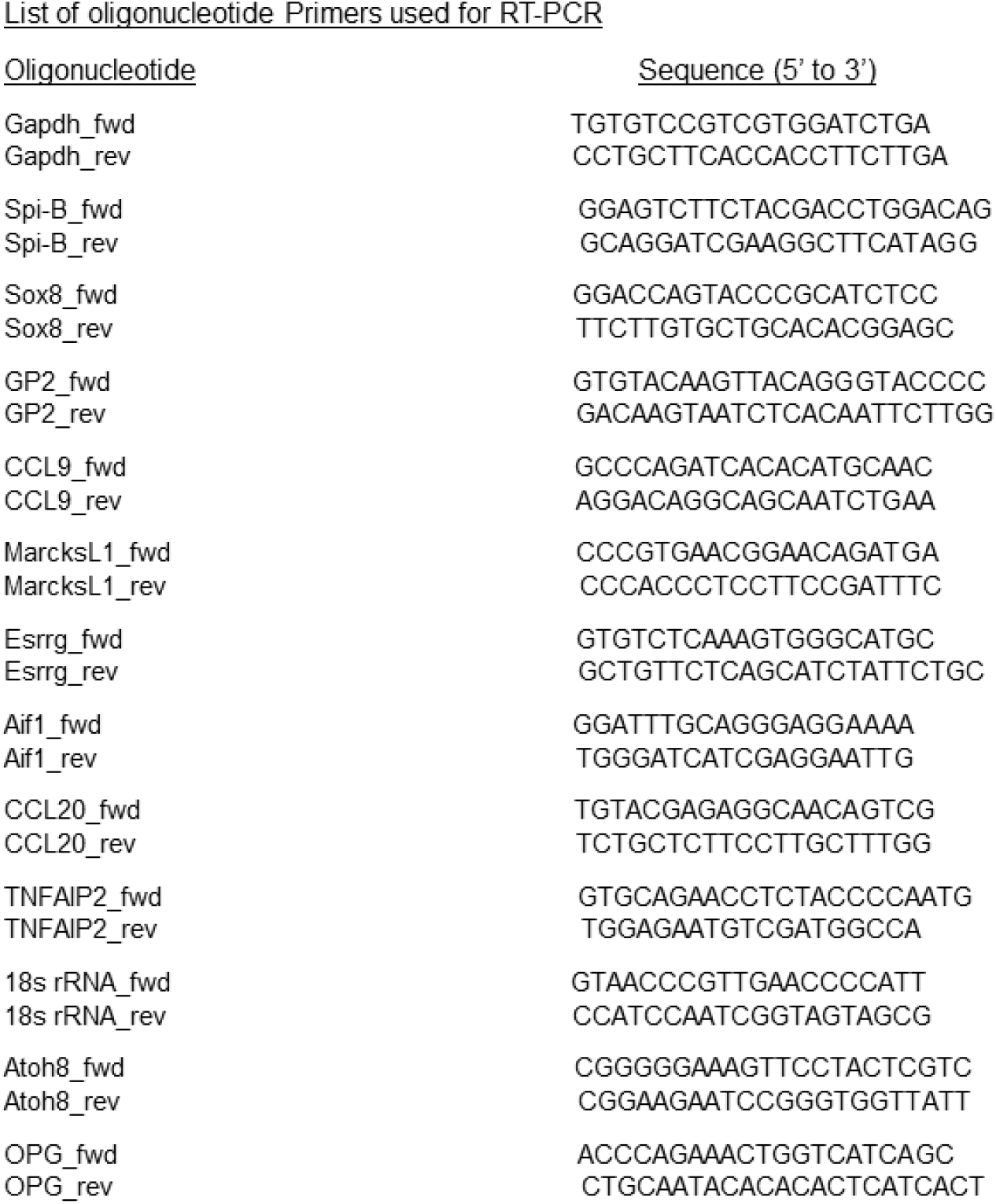
List of primers.

